# A deployable microfluidic platform for parallel and longitudinal hyperpolarised magnetic resonance metabolic phenotyping

**DOI:** 10.64898/2026.07.16.738913

**Authors:** David Gomez-Cabeza, Lluís Mangas-Florencio, Gergö Matajsz, Marc Azagra, Alba Herrero-Gomez, James Eills, Irene Marco-Rius

## Abstract

Hyperpolarised magnetic resonance (MR) enables real-time measurement of metabolic flux in living systems but remains difficult to deploy for cell-based studies because each hyperpolarisation event typically interrogates a single biological condition, limiting throughput, replication and longitudinal experimentation. Here we present a deployable microfluidic platform for parallel and longitudinal hyperpolarised ^13^C metabolic phenotyping using standard MRI instrumentation. By combining microfluidics with spatially resolved MR spectroscopic imaging, the platform converts a single hyperpolarised preparation into multiple independent metabolic measurements without dedicated radiofrequency receive arrays or specialised instrumentation. We demonstrate reproducible discrimination of metabolically active and inactive samples, resolve cell-type-specific metabolic phenotypes, quantify biochemical and pharmacological perturbations, and recover metabolic exchange kinetics from parallel samples. Beyond increasing throughput, the platform enables repeated, non-destructive metabolic interrogation of the same recirculating three-dimensional cell cultures, allowing longitudinal phenotyping of living constructs rather than endpoint comparisons of independent samples. Across this study, 186 HP-MR measurements were acquired using only 44 polarisation events, corresponding to an approximately 4-fold increase in experimental throughput, while longitudinal monitoring reduced biological sample preparation 5-fold by following the same constructs over time. By lowering the technical barrier to hyperpolarised metabolic imaging while enabling both parallel and longitudinal metabolic phenotyping, this platform provides an accessible framework for drug discovery, microphysiological systems and patient-derived models.

## 1. MAIN

Cellular metabolism is a dynamic and heterogeneous phenotype that can reflect cellular state^1,2^ environmental stress ^3,4^, disease stage and progression ^5^ and response to treatment ^6,7^. In cancer and other diseases, metabolic dysregulations often precede morphological changes and determine how cells adapt to treatments ^8^. Therefore, technologies that measure metabolism *in vivo* are paramount for drug development, disease modelling and precision medicine. Yet, due to the inherent intricacy of in vivo measurements, we still rely on in vitro ones, which only provide endpoint and indirect readouts^9^. Cell viability ^10^, extracellular metabolites ^11^ or fluorescence-based reporters ^12^ are powerful tools, but they do not directly capture the real-time conversion of metabolic substrates into downstream products. Hyperpolarised magnetic resonance (HP-MR) addresses these limitations by transiently increasing nuclear spin polarisation by several orders of magnitude, enabling real-time observation of labelled metabolic compounds^13^. Hyperpolarised [1-^13^C]pyruvate is the most widely used substrate due to its conversion into [1-^13^C]lactate, [1-^13^C]alanine and ^13^C-bicarbonate, reporting on central metabolic pathways, including glycolytic flux and amino-acid metabolism ^14–16^.

HP-MR has already shown outstanding results in preclinical ^13,17,18^ and clinical studies ^19–21^, particularly in oncology, where lactate production is a central marker of altered cancer metabolic state (i.e., Warburg effect) ^22,23^. However, HP-MR remains difficult to deploy at small scales for cellular and *ex vivo* studies.

A key challenge of HP-NMR is its throughput, because each hyperpolarised sample provides a single, short-lived signal window (i.e., seconds to minutes) that cannot be replenished and decays through the intrinsic longitudinal relaxation time constant (T_1_) of the ^13^C nuclei, the radiofrequency excitation and any metabolic or chemical conversion ^18,24,25^. As a result, one polarisation event typically interrogates one biological condition and one replicate. Because polarisation cycles require tens of minutes ^26,27^ to more than an hour ^13^ depending on the hyperpolarisation method and experimental workflow used, HP-MR experiments remain low-throughput, costly and difficult to replicate, particularly for drug-response studies, dose-response experiments and longitudinal biological models.

Several approaches have sought to address this limitation, including multi-channel receiver arrays ^28^, sequential sampling strategies ^29^ and microfluidic devices ^30^. However, each strategy has its own drawbacks. Multi-coil systems offer high sensitivity and allow high throughput, but require dedicated radiofrequency (RF) coil arrays, non-standard MRI scanners with multi-channel multi-nuclear receive capability and specialised RF engineering expertise. Microfluidic platforms reduce handling complexity but have often focused on chemical reactions ^31^ or single cell analysis ^32^, struggling to perform time-resolved measurements ^33^. Bioreactors preserve cell or tissue viability over time ^34^, but remain limited in the number of biological conditions that can be interrogated simultaneously.

We therefore asked whether the throughput and longitudinal limitations of HP-MR could be overcome by spatially encoding multiple biological samples within conventional MRI instrumentation rather than relying on specialised detector arrays. To address this, we developed a deployable microfluidic platform that combines multiwell chips with spatially resolved hyperpolarised ^13^C MR spectroscopic imaging using commercial transmit/receive volume RF coils and standard preclinical MRI acquisition methods.

Here, we show that this platform enables parallel and longitudinal HP-MR metabolic phenotyping of living cell models. Using hyperpolarised [1-^13^C]pyruvate, we simultaneously resolve multiple biological conditions within a single hyperpolarisation event, discriminate biochemical, cellular and pharmacological perturbations, recover metabolic exchange kinetics, and repeatedly interrogate the same 3D cell cultures over time. By enabling both parallelisation and longitudinal monitoring without specialised hardware, this work transforms HP-MR from a predominantly single-sample endpoint technique into an accessible platform for high-content metabolic phenotyping using MRI infrastructure already available in many preclinical imaging laboratories.

## 2. RESULTS

### 2.1. A microfluidic platform for parallel metabolic imaging

We developed a microfluidic platform to enable parallel hyperpolarised ^13^C metabolic imaging of multiple living cell samples within a single HP-MR experiment (Figure 1a-c). We produced HP [1- ^13^C]pyruvate using dissolution Dynamic Nuclear Polarisation (dDNP) (Figure 1a) and delivered through the microfluidic network to cell models contained in microfluidic chip (Figure 1b) to measure its downstream metabolism, which was detected either by chemical shift imaging (CSI) or by sequential single-pulse slice-selective (SSPSS) spectroscopy (Figure 1c) using a 3T preclinical MRI scanner. This platform allowed us to measure multiple biological samples or perturbation conditions within the same polarisation event.

**Fig. 1.**
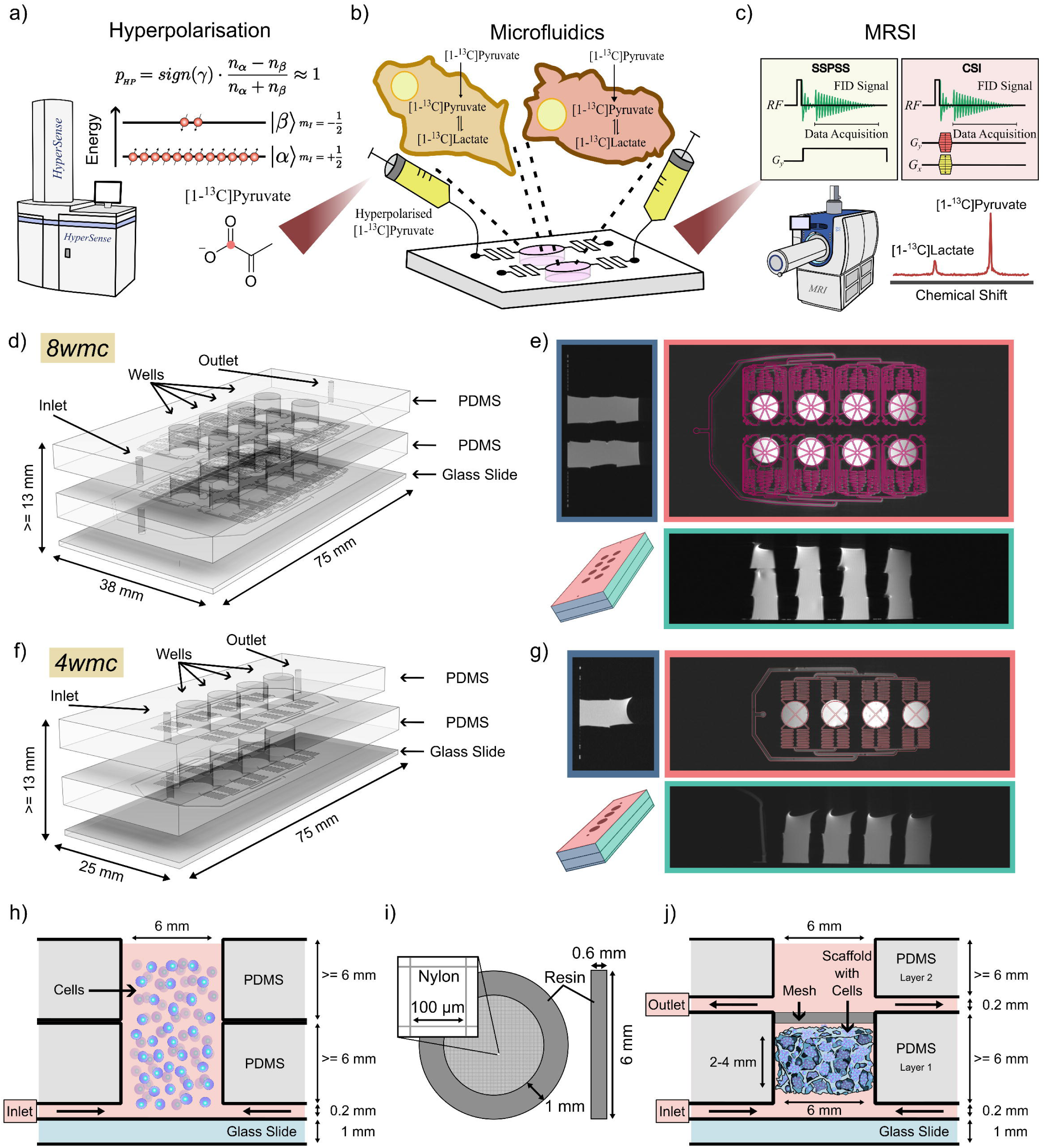
A microfluidic platform for parallel HP-MR. **a)** Dissolution dynamic nuclear polarisation (dDNP) of [1-^13^C]pyruvate for metabolic imaging. **b)** Schematic of the microfluidic platform for parallel hyperpolarised magnetic resonance (HP-MR) experiments, enabling real-time measurement of pyruvate-to-lactate conversion associated with Warburg metabolism. **c)** Magnetic resonance spectroscopy and spectroscopic imaging readouts acquired using sequential single-pulse slice selection (SSPSS) and chemical shift imaging (CSI). **d,f)** Schematics of the 8-well (8wmc) and 4-well (4wmc) microfluidic chip systems, respectively, showing chip geometry, dimensions and layer structure. **e,g)** T2-weighted MR images of the 8-well and 4-well chip systems in axial, sagittal and coronal planes. **h)** Schematic of a cell-suspension configuration within an individual well, indicating well dimensions and the position and dimensions of the inlet microfluidic channels. **i)** Schematic and dimensions of custom ring meshes designed to retain biocompatible scaffolds within the wells. **j)** Schematic of a scaffold-based 3D cell-culture configuration within an individual well, showing scaffold placement, nylon mesh retention and inlet/outlet microfluidic channels for perfusion-like substrate delivery.

The platform consists of custom multiwell (i.e., experimental chambers) microfluidic chips designed to fit within a standard preclinical MRI scanner and to operate with commercial volume RF coils. We developed two chip versions: an eight-well microfluidic chip (“8wmc”) for multiplexed spatial imaging (Figure 1d-e) and a reduced four-well (“4wmc”) one (Figure 1f-g) for experiments requiring time-resolved slice-selective acquisition. Both geometries were compatible with either cell suspensions (Figure 1h) or cell-seeded scaffolds (Figure 1j) which require a custom mesh to hold these during substrate injection and media recirculation (Figure 1i) ^35^.

The design principle was deployability. The platform does not require dedicated surface RF coils, microcoils or custom multi-channel ^13^C receiver arrays. Instead, the platform uses a commercial volume RF coil and spatial encoding to separate signals from different wells. In this way, a single short-lived hyperpolarised preparation can be converted into multiple parallel metabolic measurements while retaining compatibility with standard MRI systems.

### 2.2. Spatially resolved in-chip metabolic imaging

First, we tested whether spatially resolved imaging could discriminate chip regions containing cells in suspension from wells containing only media (i.e., control). We inserted the 8wmc in a 3D custom design microfluidic chip holder insert (Supplementary Figure 1) designed for the MRI. Then, we loaded the 8wmc with ten million human hepatoblastoma (HepG2) cells per well, and cell-free medium controls in an alternating pattern (Figure 2a). Following injection of HP [1-^13^C]pyruvate, CSIs acquired 35 s after injection (at the reaction product observable maximum) resolved substrate and [1-^13^C]lactate peaks across the field of view (Figure 2a). We detected lactate signals in wells containing HepG2 cells, whereas control wells showed negligible lactate attributed to small bleed-through (spillover) from adjacent wells, as observed when using lactate-to-pyruvate ratios (Lac/Pyr) (Figure 2b). These data define the current spatial-resolution limitation of the standard-volume-coil and CSI implementation while supporting broadly homogeneous substrate detection across the chip. SNR was comparable across all wells, with the only significant deviation observed in the upper-left well, consistent with a local edge effect. (Figure 2c). This demonstrates reproducible detection of cellular pyruvate-to-lactate conversion across multiple wells.

**Fig. 2.**
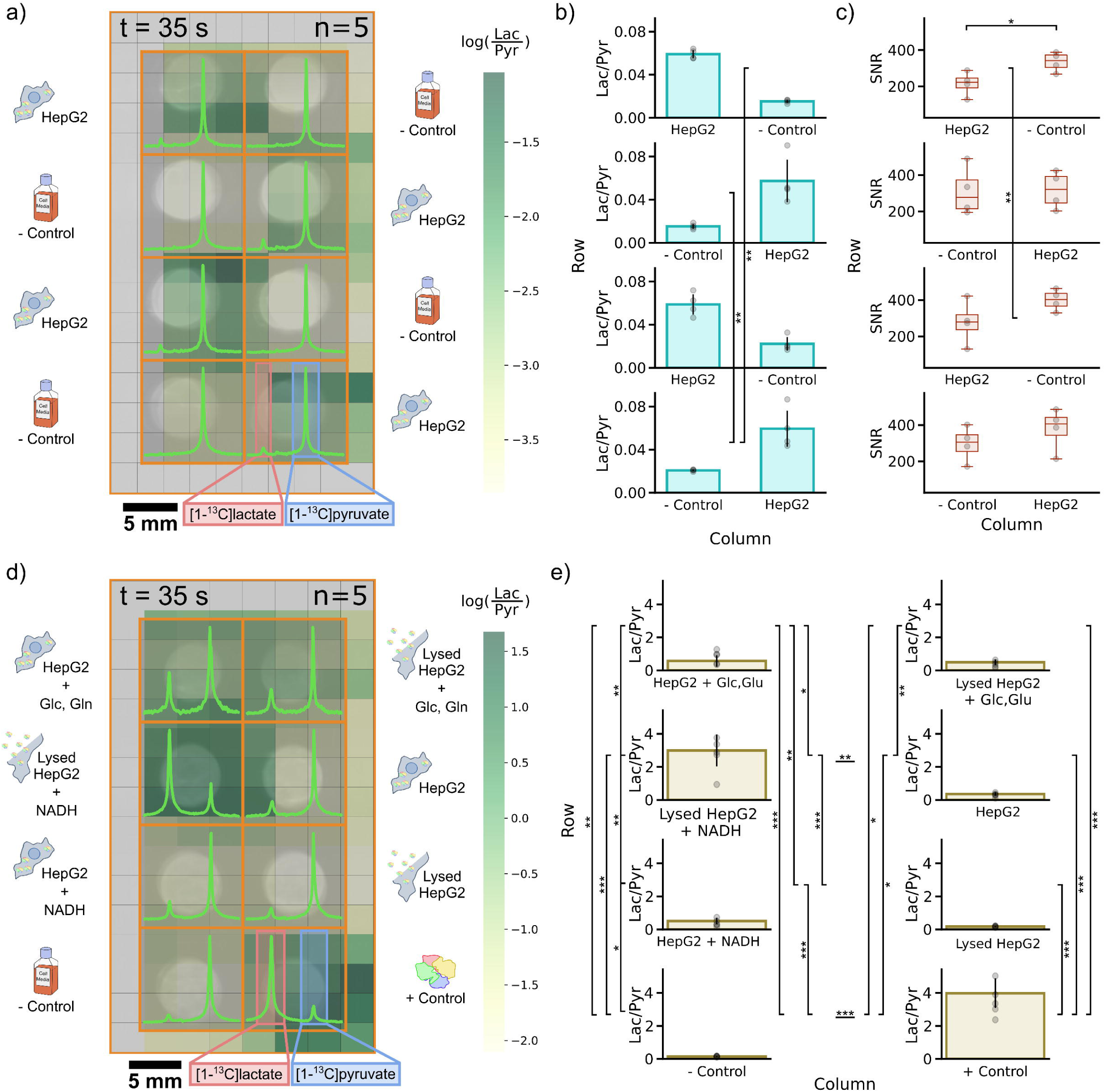
Parallel hyperpolarised magnetic resonance detects HepG2 metabolism and biochemical perturbations in an 8-well microfluidic chip. **a)** Representative chemical shift imaging (CSI) experiment in an 8-well microfluidic chip containing four wells seeded with 10 x 10^6^ HepG2 cells and four negative-control wells containing cell-free medium acquired 35 s after hyperpolarised [1-^13^C]pyruvate injection. The overlaid map shows the log-transformed [1-^13^C]lactate-to-[1-^13^C]pyruvate ratio (Lac/Pyr), calculated from peak integrals in the original 10 x 16 CSI voxel grid, on the corresponding ^1^H image of the chip. Summed ^13^C spectra from all voxels assigned to each well are shown within the corresponding well regions. Voxels containing only noise are displayed in white for visualisation purposes; this convention is used for all CSI grids shown throughout the manuscript. **b)** Lac/Pyr measured across four independent replicates of the experiment shown in **a**. HepG2-containing wells showed significantly higher Lac/Pyr than negative-control wells, whereas no significant differences were observed within each group, except for the indicated pairwise comparisons. **c)** [1-^13^C]pyruvate SNR quantified for each well region in the experiments shown in **a**, used to assess position-dependent variation in pyruvate detection across the chip. **d)** Representative CSI experiment in an 8-well microfluidic chip containing six HepG2 experimental conditions, each with 10 x 10^6^ cells per well, plus one negative-control well containing medium only and one positive-control well containing commercial rabbit lactate dehydrogenase (LDH). Conditions comprised intact or lysed HepG2 cells in standard medium, medium supplemented with glucose and glutamine (+Glc/Gln) or medium supplemented with NADH. The log-transformed Lac/Pyr map and summed spectra are displayed as in **a**. **e)** Lac/Pyr measured across five independent replicates of the experiment shown in **d**, showing condition-dependent differences within a single chip. Data are shown as mean ± SD. Statistical significance is indicated as *0.05 ≥ p > 0.01, **0.01 ≥ p > 0.001, *** p ≤ 0.001.

### 2.3. Multiplexed biochemical perturbations

Having established spatially resolved cell-versus-control discrimination, we also validated that our platform could resolve defined biochemical perturbations affecting Lac/Pyr within a single experiment. We loaded the 8wmc with intact HepG2 cells, lysed HepG2 cell extracts, and corresponding conditions supplemented with either NADH or glucose and glutamine to probe the influence of cofactor availability and substrate metabolism on pyruvate-to-lactate conversion. Cell-free medium and purified LDH served as negative and positive controls, respectively, resulting in eight biochemical conditions analysed simultaneously (Figure 2d-e).

The metabolic maps obtained showed condition-specific differences across the 8wmc (Figure 2d). Supplementation of intact HepG2 cells with glucose and glutamine increased Lac/Pyr by an average of 163% relative to untreated cells (Figure 2e), consistent with enhanced glycolytic activity ^36^. Addition of extracellular NADH produced a more modest increase (32%), due to the role of extracellular NADH in adjusting the redox balance ^37,38^. In contrast, lysed cell extracts exhibited substantially lower Lac/Pyr than intact cells, due to disturbed redox balance and decrease in LDH concentration of the sample (∼0.28 times). Supplementation of lysates with glucose and glutamine increased Lac/Pyr by 117%, whereas NADH supplementation produced a striking 16-fold increase over untreated lysates, indicating that cofactor availability becomes the dominant limiting factor once membrane transport barriers are removed (Rao et al., 2020). As expected, purified LDH with excess NADH generated the highest Lac/Pyr, whereas cell-free medium showed negligible lactate production.

Collectively, these conditions establish the biochemical specificity of the HP-MR readout. The progressive increase in Lac/Pyr from cell-free controls to intact cells, metabolically stimulated cells, lysates supplemented with NADH and purified enzyme demonstrates that the HP lactate signal is governed by enzymatic pyruvate-to-lactate exchange and modulated by substrate availability, cofactor supply and cellular compartmentalisation, rather than by pyruvate delivery or non-specific chemical conversion alone. Importantly, because all conditions were measured simultaneously, comparisons were performed under identical polarisation, injection, temperature and acquisition conditions, eliminating a major source of variability in conventional HP-MR experiments.

### 2.4. Parallel imaging resolves cell-type-specific metabolic phenotypes

We next tested whether the platform could resolve metabolic differences between distinct cell types within a single hyperpolarised experiment. We loaded HepG2 and human adenocarcinoma (HeLa) cells into alternating wells of the 4wmc and imaged them through CSI after delivery of HP [1-^13^C]pyruvate (Figure 3a).

**Fig. 3.**
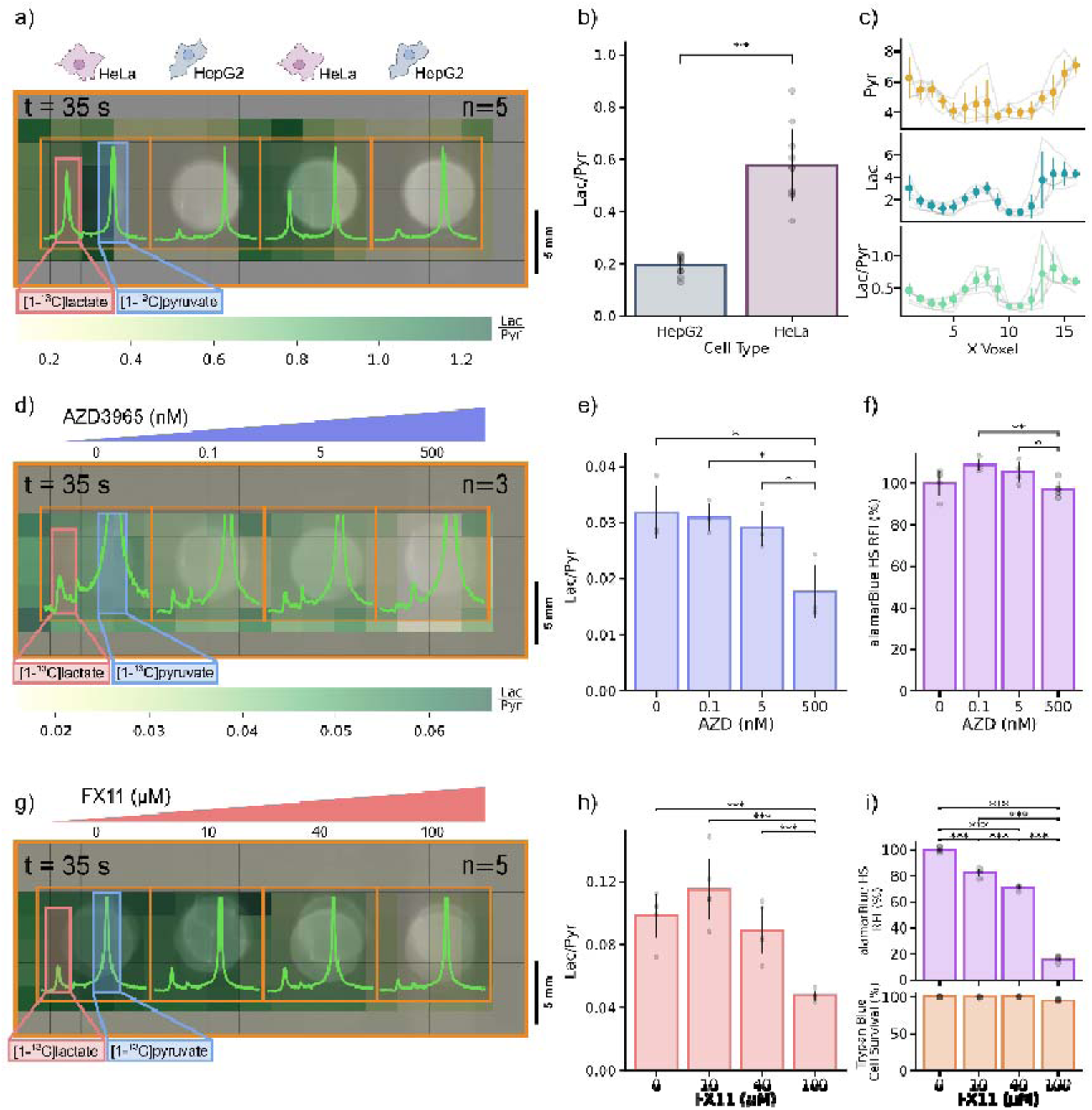
Parallel metabolic phenotyping resolves cell identity and drug response. Representative chemical shift imaging (CSI) experiments in the 4-well microfluidic chip comparing: **a**) two cell types HeLa and HepG2 cells, and **d**) HepG2 cells exposed to increasing concentrations of the MCT1 inhibitor AZD3965 or **g**) the LDH inhibitor FX11. The overlaid maps show the [1-^13^C]lactate-to-[1-^13^C]pyruvate peak-integral ratio (Lac/Pyr), calculated from the original 8 x 16 CSI voxel grids, on the corresponding ^1^H anatomical images of the chip. Summed ^13^C spectra from all voxels assigned to each well are shown overlaid on top of the corresponding image. Voxels containing only noise are displayed in white for visualisation purposes and were excluded from spectral summation; this convention is used for all CSI grids shown throughout the manuscript. **b,e,h)** Quantification of Lac/Pyr for each condition in the experiments shown in **a**, **d** and **g**, respectively, across five independent replicates for the cell-type comparison (**b**) and three independent replicates for each drug experiment (**e,h**). **c)** Voxel-wise profiles of pyruvate signal intensity (top), lactate signal intensity (center) and Lac/Pyr (bottom), summed along the y-axis of the chip shown in **a**. Points show the mean voxel intensity across replicates, error bars indicate s.d., and grey lines show individual experimental replicates. **f,i)** HepG2 cell viability measured after ≈ 30 min exposure to AZD3965 (**f**) or approximately 24 h exposure to FX11 (**i**), before the hyperpolarised experiments shown in **d** and **g**, respectively. Data are shown

HeLa cells produced substantially higher Lac/Pyr values than HepG2 ones across biological replicates (Figure 3b). Voxel-wise profiles confirmed that lactate signal and Lac/Pyr co-localised with HeLa-containing wells, whereas pyruvate signal remained comparable across wells (Figure 3c). Thus, the observed differences were not primarily driven by uneven substrate delivery or acquisition bias, but by cell-type-dependent metabolism.

Measuring both cell types within the same polarisation event reduces confounding from run-to-run differences in polarisation level, injection timing and acquisition conditions. This internally controlled format is relevant for comparing isogenic models under different cellular environment conditions, patient-derived samples or treatment testing.

### 2.5. Pharmacological lactate modulation detection through MRSI

Our platform further quantified drug effects in the LDH pathway. Here, we used two inhibitors acting at different points in the pathway: 1) AZD3965, an inhibitor for the pyruvate monocarboxylate transporter 1 (MCT1) ^40,41^, and 2) FX11, which inhibits LDH activity ^42,43^.

HepG2 cells treated with AZD3965 showed a dose-dependent reduction in Lac/Pyr (Figure 3d-e). At the highest drug concentration (500 µM), Lac/Pyr decreased by 58±20% relative to non-treated cells. This occurred without a corresponding reduction in cell viability (Figure 3f), validating a pharmacodynamic effect on pyruvate transport. As expected, lower drug concentrations resulted in small lactate production reductions.

FX11 also reduced lactate production in a dose-dependent manner (Figure 3g,h). At low drug concentrations (0-40 µM), lactate production showed small changes, despite a progressive decrease cell viability (Figure 3i) but no detectable change in cell survival (Figure 3j). At 100 µM, lactate production decreased by 49±8%, accompanied by a drop of ∼84±2% reduction cell viability (Figure 3i). These data proved that high-dose FX11 suppresses LDH activity. These experiments show that our platform detects pathway-specific metabolic perturbations in parallel and the combination of HP-MRI with viability assays helps distinguish pharmacodynamic metabolic inhibition from cytotoxicity.

### 2.6. Parallel time-resolved slice-selective acquisition

The previous experiments used single-time-point CSI to compare Lac/Pyr across multiple wells. We next tested whether the platform could also recover dynamic metabolic information in parallel implementing a sequential single-pulse slice-selective (SSPSS) spectroscopy sequence. Using the 4mwc, we acquired time-resolved ^13^C spectra from three wells containing HepG2 cells and one cell-free control one after injection of HP [1-^13^C]pyruvate (Figure 4a,b). In HepG2-containing wells, pyruvate signal increased after substrate arrival and then decayed due to longitudinal relaxation, metabolic conversion and RF excitation. The lactate signal appeared after pyruvate delivery (Figure 4c). In contrast, the control well showed negligible lactate spillover signal, confirming cellular metabolism. We confirmed the results by observing a high Lac/Pyr ratio in wells containing cells and a negligible one in the control one (Figure 4d).

**Fig. 4.**
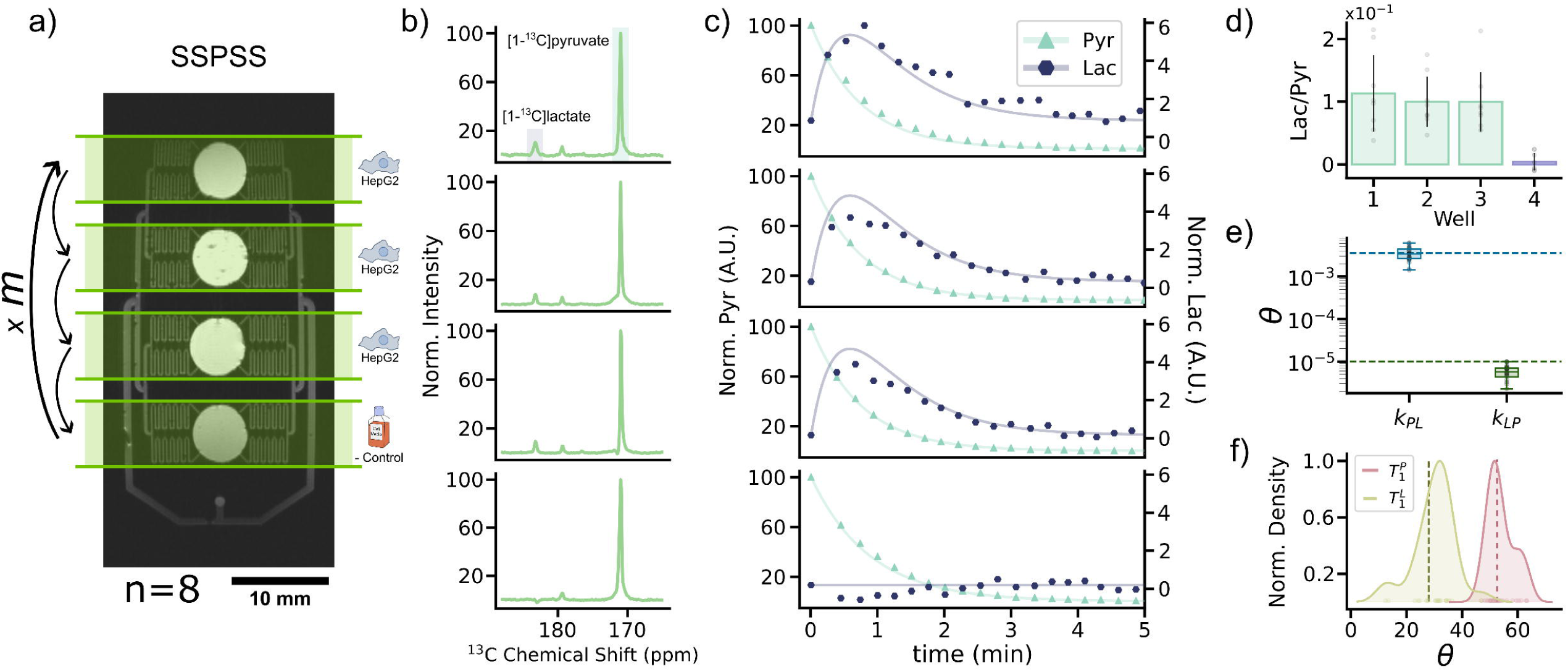
Time-resolved parallel dynamic hyperpolarised metabolic imaging enables kinetic analysis in microfluidic chips. **a)** Schematic of the sequential single-pulse acquisition strategy used in a 4-well microfluidic chip (4wmc). Acquisition proceeded from the top well to the bottom well sequentially and as repeated for *n* cycles. **b)** Time-summed ^13^C spectra from each well in a representative sequential 4wmc experiment. **c)** Representative time courses of [1-^13^C]pyruvate and [1-^13^C]lactate signals acquired from each well in the experiment shown in **b**. As indicated in **a**, the top three wells contained 10 x 10^6^ HepG2 cells per well, whereas the bottom well contained cell-free medium. Lines show model simulations obtained after estimating a single set of kinetic rate constants and *T*_1_ longitudinal relaxation times using all data acquired in this study. **d)** Lactate-to-pyruvate (Lac/Pyr) area-under-the-curve ratio for each acquired well. Bars show mean ± SD. across independent replicates, and overlaid points show individual replicate values. **e)** Estimated pyruvate-to-lactate (*k*_PL_) and lactate-to-pyruvate (*k*_LP_) kinetic rate constants. **f**) Estimated pyruvate (*T*_1_^P^) and lactate (*T*_1_^L^) longitudinal relaxation time constants obtained by fitting each experiment independently. Dashed lines indicate the global parameter estimates obtained by fitting all experiments simultaneously.

We fitted the pyruvate and lactate data using a two-site exchange model (see Supplementary Material) incorporating forward pyruvate-to-lactate conversion (*k*_PL_), reverse exchange (*k*_LP_) and apparent relaxation (*T*_1_). The fitted *k*_PL_ was consistently higher than *k*_LP,_ as expected for net LDH activity in HepG2 cells (Figure 4e). The model also recovered distinct apparent *T*_1_ longitudinal relaxation behaviour for pyruvate and lactate, with lactate showing an ∼2 -fold shorter detection window than pyruvate (Figure 4f). These data show that the platform is not restricted to endpoint metabolite ratios. When temporal sampling is available, it can support kinetic modelling directly within the microfluidic chip format.

### 2.7. Longitudinal HP metabolic monitoring of cell-laden scaffolds

Finally, we demonstrated that the platform supports repeated HP-MR measurements of scaffold-based 3D cell cultures, allowing longitudinal metabolic monitoring. We seeded HepG2 cells into carboxymethylcellulose (CMC) scaffolds (Supplementary Figure 2), and loaded them into the 8wmc, connecting it to a closed media recirculation circuit (Supplementary Figures 3-4). Between imaging sessions, we incubated the chip with re-circulating media, repeating up to five HP [1-^13^C]pyruvate injections (Figure 5a-b). Due to the low number of cells enclosed in the scaffold after seeding (Supplementary Figure 5), we performed a single acquisition of two groups of four wells to assess lactate production (Figure 5c).

**Fig. 5.**
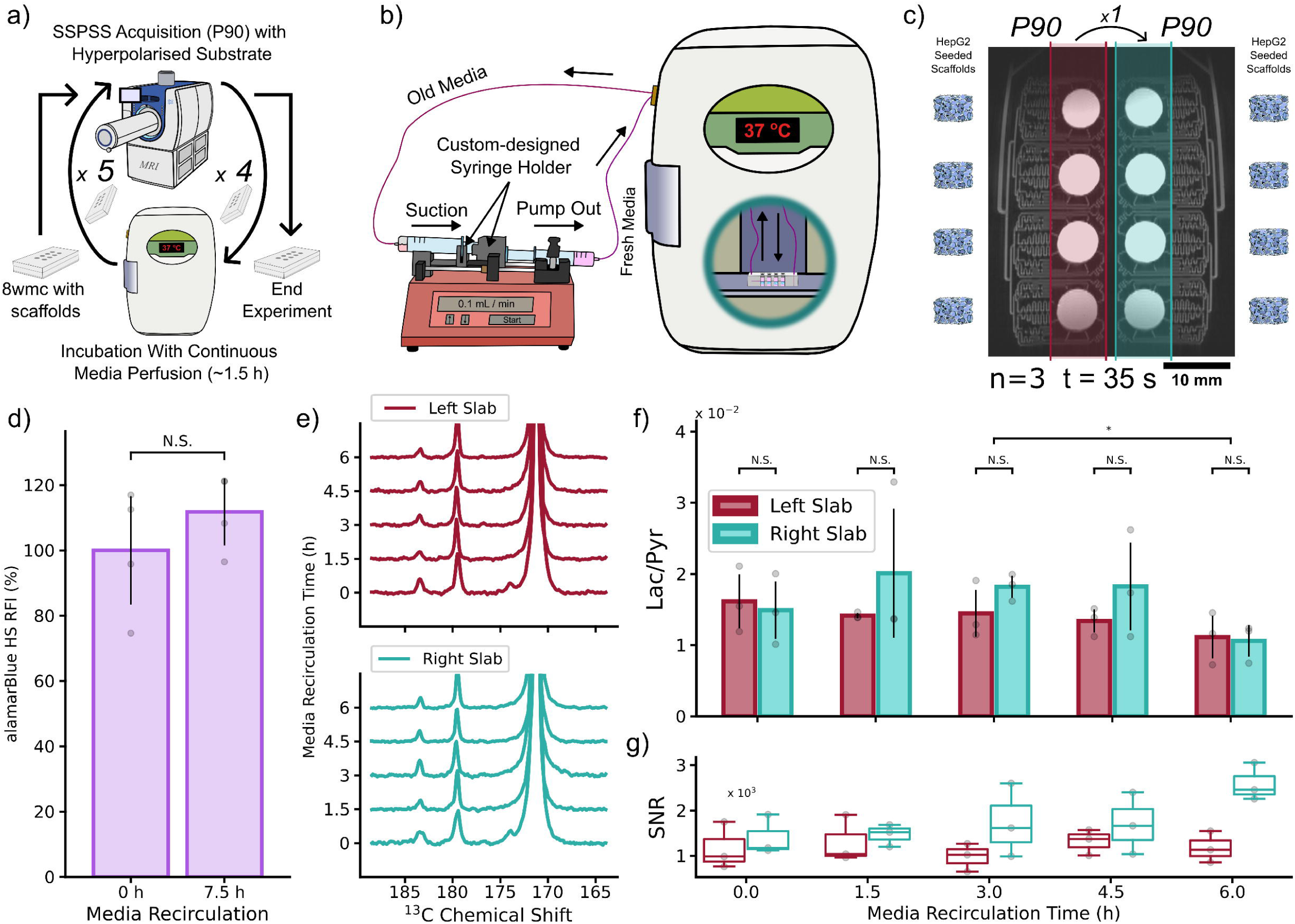
Longitudinal parallel hyperpolarised metabolic imaging monitors metabolism over time in perfused 3D HepG2-ladden scaffolds. **a)** Workflow for longitudinal hyperpolarised metabolic imaging experiments. **b)** Schematic of the medium-recirculation system used during the incubation period. **c)** Acquisition protocol for repeated hyperpolarised measurements over the longitudinal experiment. **d)** AlamarBlue high-sensitivity (HS) relative fluorescence intensity (RFI) of HepG2-seeded scaffolds maintained under static conditions (0 h) or after five rounds of medium recirculation in the 8-well microfluidic chip (8wmc) over 7.5 h. **e)** Representative stacked ^13^C spectra acquired after each medium-recirculation period from two sampled chip regions containing HepG2-seeded CMC scaffolds: left region shown in red and right region shown in blue. **f)** Lactate-to-pyruvate ratio (Lac/Pyr) measured in each sampled region after increasing cumulative medium-recirculation times (n=4). **g)** Signal-to-noise ratio (SNR) for the experiments shown in **f**. Data are shown as mean ± SD. Statistical significance is indicated as *0.05 ≥ p > 0.01, **0.01 ≥ p > 0.001, *** p < 0.001.

The total cell content within each scaffold was assessed using metabolic activity and total protein assays (Supplementary Figure 5). Although the porous cryogel had validated flow-in properties compatible with HP-MR experiments and supported cell culturing ^35^, cell seeding remained technically challenging, allowing a maximum of ∼3 x 10^6^ cells per scaffold. Several seeding protocols were evaluated with the most reproducible approach consisting of 4 repeated seedings of two millions cells on two consecutive days (two seedings per day) into each scaffold. We confirmed that scaffolds maintained cell viability during repeated media recirculation for up to 7.5 hours (Figure 5d).

HP-MR spectra acquired after each 1.5 h recirculation consistently resolved [1-^13^C]pyruvate and [1-^13^C]lactate peaks in both sampled chip regions (Figure 5e). Lac/Pyr ratios remained largely stable over the 7.5 h monitoring period, with only a modest decline observed at the final measurement, most likely reflecting local variation in substrate access or perfusion efficiency rather than a metabolic temporal change (Figure 5f). We also obtained the Lac/Pyr ratios evolution of each chip individually as a function of incubation time (Supplementary Figure 6).

No significant differences were observed between the two acquisition regions, further demonstrating the reproducibility of the platform across independent sampling locations. Likewise, SNR remained stable throughout the experiment (Figure 5g), indicating that repeated measurements did not compromise data quality.

These results demonstrate that the proposed platform supports repeated HP-MR metabolic measurements in 3D cell-seeded scaffolds under recirculating culture conditions using conventional MRI hardware. This establishes a practical bridge between suspension-cell HP-MR assays and longitudinal metabolic monitoring in more physiologically relevant microfluidic 3D cell culture systems.

## 3. DISCUSSION

We developed a deployable microfluidic platform for parallel and longitudinal hyperpolarised ^13^C metabolic imaging of living cell models using standard commercial MRI equipment. The main advantage of our platform is its architecture, as it increases the accessibility, reproducibility and longitudinal compatibility of cell-based HP-MRI studies. By combining microfluidics, spatially resolved spectroscopic imaging and commercial RF coils, the platform enables measurement of several independent biological samples or conditions within a single HP experiment, without dedicated ^13^C receive arrays or custom multi-channel RF hardware (Figure 1).

Our design addresses a major practical limitation of cell-based HP-MR: throughput. Hyperpolarised signals are short-lived and non-renewable, such that conventional workflows often use one polarisation event to interrogate one sample,one replicate of one condition or time point. This hinders reproducibility and makes multi-condition studies expensive and time-consuming ^44^. Spatially encoding of multiple interrogation wells converts a single HP experiment into several internally controlled measurements acquired under matched polarisation, injection and acquisition conditions. Across this study, 186 HP-MR measurements were acquired using only 44 dDNP events, corresponding to an approximately 4.2-fold increase in throughput compared with a conventional one-condition-per-polarisation workflow. Assuming a typical [1-^13^C]pyruvate dDNP polarisation time of 1 h and based on the published institutional service rates for MRI scanner time, dDNP and technical support at the Institute for Bioengineering of Catalonia (https://ibecbarcelona.eu/services/price-list/), this corresponds to 142 h of polarisation time saved and an estimated cost reduction of approximately €28,400 (Supplementary Table 1).

The longitudinal capability provides a second, conceptually distinct advantage. Conventional endpoint designs require independent biological samples for each time point. In contrast, our platform repeatedly interrogates the same living 3D cultures over time. In the longitudinal experiments presented here, 3 scaffold cultures generated measurements across 5 time points, whereas a conventional endpoint design would have required 15 independently prepared scaffolds. This 5-fold reduction in biological sample preparation reduces workload, decreases inter-sample variability and enables true repeated-measures experimental designs, such as drug-response studies and comparative metabolic phenotyping. These advantages are particularly relevant for organ-on-chip systems, patient-derived cultures and other scarce, expensive or labour-intensive biological models.

Our biochemical control experiments define the mechanistic basis of the lactate-to-pyruvate readout. HP [1-^13^C]lactate production is often interpreted as a marker of glycolytic activity, but the measured signal reflects multiple coupled processes, including pyruvate delivery, membrane transport, LDH activity, NADH availability, cellular compartmentalisation or relaxation and acquisition timing ^45–47^. By combining intact cells, lysed cells, NADH-supplemented conditions, glucose- and glutamine-supplemented conditions, purified LDH and cell-free controls within the same chip, we showed that pyruvate-to-lactate conversion followed a graded biochemical response (Figure 2). The increase in lactate production after glucose and glutamine supplementation is consistent with enhanced glycolytic activity ^36^, whereas the response to NADH confirms the contribution of redox state and cofactor availability ^37^. The strong lactate production in NADH-supplemented lysates and purified enzyme controls further supports the enzymatic origin of the signal. These data show that Lac/Pyr reports LDH-mediated pyruvate-to-lactate exchange modulated by substrate access, cofactor supply and cellular organisation, rather than non-specific conversion or signal mis-registration.

The cell-line and inhibitor experiments show that this readout can be used for functional metabolic phenotyping (Figure 3). HeLa cells showed higher Lac/Pyr than HepG2 cells despite comparable pyruvate delivery, consistent with cell-type-dependent differences in pyruvate-to-lactate exchange ^48,49^. Because both cell lines were measured within the same hyperpolarisation experiment, the comparison is less confounded by inter-experiment variation in polarisation level or substrate concentration, injection timing or scanner conditions. AZD3965, an MCT1 inhibitor currently in clinical trials ^50^, reduced Lac/Pyr without reducing alamarBlue signal, consistent with inhibition of MCT-mediated pyruvate uptake before overt loss of cellular reducing activity ^41^. FX11 decreased Lac/Pyr at high concentration, accompanied by reduced alamarBlue signal, while trypan blue exclusion remained unchanged. These results indicates suppression of pyruvate-to-lactate conversion by direct inhibition of LDH enzyme and a reduced cellular metabolic activity without affecting cell survival of attached cells treated with the inhibitor ^51,52^. Combined, these experiments illustrate the ability of HP-MR to detect pathway disruption in real-time and in intact cells, providing information not always accessible through conventional endpoint assays alone.

The time-resolved experiments extend the platform usability beyond static Lac/Pyr mapping. Sequential single-pulse slice-selective spectroscopy measured pyruvate and lactate dynamics across spatially separated wells, and two-site exchange modelling recovered apparent exchange and relaxation parameters (Figure 4). This distinction is important because metabolite ratios and kinetic parameters are not equivalent. Lac/Pyr provides a robust, high-throughput phenotyping metric, whereas kinetic modelling can help separate contributions from substrate transport, enzyme activity, redox state and relaxation when sufficient temporal information is available ^53,54^. Although the current implementation was not optimised for maximal kinetic precision, it shows that dynamic metabolic modelling can be incorporated into the microfluidic workflow without specialised detector hardware.

The longitudinal 3D experiments move the platform towards more representative non-destructive metabolic monitoring ^55,56^. Most cell-based HP-MR workflows remain endpoint measurements in suspensions, tissue slices or single-use preparations ^57–59^. Here, the same cell-seeded scaffolds were measured repeatedly by HP-MR over 6 h under media recirculation, while maintaining alamarBlue signal, Lac/Pyr stability and SNR (Figure 5). The alamarBlue data should be interpreted as preservation of cellular reducing activity, rather than definitive proof of unchanged cell number or viability. Nevertheless, stable HP-MR readouts and retained signal quality show that repeated metabolic measurements are feasible in a recirculating 3D microfluidic configuration.

HP-MR of cell-laden scaffolds depends on the biomaterial properties. HP-MR requires rapid and homogeneous delivery of the hyperpolarised substrate because the signal decays within seconds to minutes. The highly porous cryogel scaffold used here allowed hyperpolarised pyruvate to reach cells throughout the construct within seconds of injection ^35,60^, making it compatible with the HP-MR time window. Not all biomaterials used to provide 3D structure to cell cultures will meet this requirement. An additional advantage of this scaffold is that cell-seeded constructs can be cryopreserved and re-analysed at later time points, which is relevant for scarce or valuable samples ^61^. The current model remains intentionally simple: it uses a single cancer cell line and does not reproduce the multicellular architecture, barrier function, oxygen gradients or mechanical complexity of mature organ-on-chip systems. However, it establishes the technical basis for integrating HP-MR with more advanced microphysiological models.

The main technical limitation of the current implementation is sensitivity. Commercial volume RF coils provide straightforward implementation, compatibility with standard MRI scanners and a broad and homogeneous B_0_ field coverage, but they offer lower local sensitivity than surface coils, microcoils or dedicated multi-channel receive arrays ^28^. This increases the minimum number of cells required per experiment, limits spatial resolution and contributes to residual voxel spillover between neighbouring wells. This trade-off was intentional. The objective was not to maximise SNR, but to demonstrate that useful parallel and longitudinal HP-MR measurements can be achieved with infrastructure already available in many imaging laboratories. In future implementations, the microfluidic platform could be scaled further by tailoring well number and geometry to the metabolic activity of the cell model, and by integrating cross-talk channels to modulate metabolism between compartments. Furthermore, for applications involving scarce samples or low cell numbers, future implementations could incorporate local receive arrays with improved coil geometries or multi-channel multinuclear architectures.

Several biological and acquisition limitations remain. Established cancer cell lines and single-cell-type scaffolds provide reproducible metabolic responses for platform validation, but they do not capture the heterogeneity of primary tissues, organoids or organ-on-chip systems. Translation to more physiological models will require optimisation of scaffold architecture, perfusion, oxygenation, extracellular matrix composition, substrate delivery and normalisation strategies. Acquisition efficiency can also be improved. The CSI and slice-selective methods used here prioritise robustness and compatibility with standard scanners, but faster spectroscopic imaging, compressed sensing, variable flip-angle schemes and spectral-spatial excitation could improve temporal resolution and preserve hyperpolarised magnetisation more efficiently. These developments will be important for lower-cell-number models and for detecting lower-abundance metabolites such as alanine and bicarbonate.

Overall, this work shifts cell-based HP-MR from a predominantly single-sample endpoint assay towards a parallel and longitudinal metabolic phenotyping platform. By combining standard MRI hardware with spatially encoded microfluidics, the platform increases throughput, reduces experimental cost, enables repeated measurements of the same living 3D cultures and preserves mechanistic metabolic readouts. This provides a practical foundation for integrating HP-MR with drug-response studies, organ-on-chip technologies, patient-derived models and translational metabolic screening workflows.

## 4. METHODS

### 4.1. Cell culture

HepG2 human hepatoblastoma cells (CliniSciences) and HeLa human adenocarcinoma cells (ATCC CCL-2) were cultured in Eagle’s Minimum Essential Medium (EMEM; ATCC, 30-2003) supplemented with 10% fetal bovine serum (FBS) and 1% penicillin–streptomycin (P/S) at 37°C in a humidified atmosphere containing 5% CO_2_. Culture medium was replaced every 48-72 h as required.

For experiments, cells were used at ≥ 80% confluence. Cells were washed with phosphate-buffered saline (PBS) and detached using 0.25% trypsin-EDTA. Cell suspensions were centrifuged at 200g for 3 min and resuspended in 1 mL of culture medium for cell counting using an automated cell counter (CountessTM 3.0, Thermo Fisher Scientific). Cells were then centrifuged again and resuspended at the desired concentration and final volume required for each microfluidic experiment.

### 4.2. Additional incubations and drug treatments

Where indicated, culture medium was supplemented with glucose and glutamine at final concentrations of 25 mM and 4 mM, respectively. For experiments requiring exogenous reaction cofactor supplementation, NADH was added at a final concentration of 33 mM. Cell lysates were prepared using RIPA lysis buffer when specified.

For drug treatment experiments, HepG2 cells were exposed to FX11 (427218, Merck), an LDH inhibitor, or AZD3965 (HY-12750, MedChemExpress), an MCT1 inhibitor. FX11 treatments were performed at 0 μM (0.1% DMSO vehicle control), 10 μM, 40 μM or 100 μM for 24 h. AZD3965 treatments were performed at 0 nM (0.1% DMSO vehicle control), 0.1 nM, 5 nM or 500 nM for 20 min.

Cell viability was assessed using trypan blue staining with an automated cell counter (CountessTM 3.0, Thermo Fisher Scientific) and with the alamarBlue HS assay (Thermo Fisher Scientific). Fluorescence measurements were acquired using a microplate reader (Synergy H1, BioTek) with excitation/emission wavelengths of 560/590 nm.

### 4.3. CMC scaffold fabrication

Carboxymethyl cellulose (CMC) cryogel scaffolds were fabricated using previously reported methodologies (10.1088/1758-5090/ac00c3). Briefly, a 1% (w/v) solution of 90 kDa CMC was prepared in Milli-Q water under constant stirring. A crosslinking solution composed of 50 mM 2-(N-morpholino)ethanesulfonic acid (MES) buffer adjusted to pH 5.5 with NaOH, adipic acid dihydrazide (ADH; 1.83 mM) and N-(3-dimethylaminopropyl)-N′-ethylcarbodiimide hydrochloride (EDC; 18.9 μM) was added to the solution.

To minimise premature crosslinking, the solution was continuously stirred during dispensing into polydimethylsiloxane (PDMS) moulds consisting of annular cylinders (4 mm height, 10 mm diameter) positioned on 24 x 24 mm glass coverslips. Samples were frozen overnight at -20 °C and thawed the following day. Cryogels were subsequently cut into 6 mm diameter discs using a biopsy punch, immersed in PBS and sterilised by autoclaving.

### 4.4. Cell seeding in biocompatible cryogels

CMC cryogel scaffolds were placed into 6-well plates containing PDMS retaining rings to stabilise the scaffolds during seeding. Cell seeding was performed over two consecutive days.

On day 1, 4 x 10^6^ cells were seeded onto each scaffold in two sequential steps consisting of an initial 2 x 10^6^ cell suspension followed by an additional 2 x 10^6^ cells after 20 min. On day 2, two further seeding steps of 2 x 10^6^ cells each were performed to achieve a final theoretical density of 8 x 10^6^ cells per scaffold. (Supplementary Figure 2).

After 24 h, scaffolds were transferred to fresh culture plates and medium was replaced every 48 h thereafter. To reduce biological variability of the scaffolded cells, we used the scaffolds between 24-48h after the second seeding.

### 4.5. Microfluidic chip fabrication

Two versions of the microfluidic platform were designed using CleWin 4.1 and Autodesk Fusion to accommodate different experimental throughput requirements (Figure 1d-g). The devices contained either four or eight experimental wells within footprints of 75 x 25 mm and 75 x 38 mm, respectively (Figure 1d,f).

The microfluidic layouts incorporated a substrate injection inlet connected to a network of channels with a constant height of 200 μm and widths decreasing from 800 μm to 200 μm. At each bifurcation, channels were designed with equal path lengths to promote homogeneous flow distribution^30^. The channels terminated in circular outlet geometries distributed equidistantly around the 6 mm diameter cylindrical experimental wells (Figure 1h) to facilitate homogeneous substrate perfusion.

Devices were fabricated using standard soft lithography techniques from 5 ± 1 mm thick polydimethylsiloxane (PDMS; Sylgard 184, Sigma-Aldrich) bonded to 1 mm thick glass slides (Corning).

### 4.6. Nylon mesh fabrication

Scaffold support meshes for the microfluidic chips (Figure 1i) were fabricated as described previously^35^. Briefly, meshes were fabricated using a digital light processing (DLP) 3D printer (Versus 385 nm, Microlay, Spain). Biocompatible resin frames (SolusArt v3.0, Gesswein, USA) were printed directly onto 100 μm pore-size nylon mesh sheets. For fabrication, nylon mesh sheets were secured to the printer build platform using double-sided adhesive tape (No. 99786, 3M, USA). Ring-shaped resin frames with a height of 0.6 mm, an outer diameter of 6 mm and an inner diameter of 5 mm were subsequently printed and cured directly onto the mesh.

### 4.7. [1-^13^C]pyruvate hyperpolarisation using dissolution DNP

Dissolution dynamic nuclear polarisation (dDNP) was used to hyperpolarise [1-^13^C]pyruvate for all experiments. Cell suspension studies were performed using a HyperSense polariser (Oxford Instruments Ltd., UK), whereas scaffold-based experiments were performed using a SpinAligner system (Polarize, Denmark).

HyperSense stock samples consisted of [1-^13^C]pyruvic acid (14.2 M, Merck Life Sciences), OX063 radical (18.1 mM, GE Healthcare) and DOTA (1.4 mM, Guerbet). SpinAligner samples consisted of [1-^13^C]pyruvic acid (14.4 M, Merck Life Sciences) and AH radical (30.0 mM, Polarize).

Following polarisation, samples were dissolved using a heated dissolution buffer containing HEPES (40.0 mM), NaOH (94.0 mM), EDTA-H4 (0.3 mM) and NaCl (3.0 mM) in Milli-Q water to obtain a final [1-13C]pyruvate concentration of 40 mM.

### 4.8. Chip- and syringe pump holder design and fabrication

Custom-made supporting equipment (e.g., chip inserts, syringe holders) were designed using Fusion (Autodesk, USA). The parts were 3D printed via fused deposition modelling (FDM) from polylactic acid (PLA) filaments (RS Components, UK) using either an Ender-3 S1 Pro (Creality, China) or a Bambu Lab H2S (Bambu Lab, China) printer. Schematics shown in Supplementary Figure 1 and 3.

### 4.9. Parallel HP-MR analysis of cell-laden microfluidic platforms

A schematic overview of the complete platform is shown in Figure 1. Experiments were performed using either a four-well (4wmc) or eight-well (8wmc) microfluidic chip. Microfluidic devices were mounted onto the animal handling system of a Bruker BioSpec 3 T MRI scanner (Bruker, Germany). PTFE tubing was used to reinforce microfluidic outlet connections, while nylon tubing was used for delivery of hyperpolarised substrates into the experimental wells. Polypropylene male Luer-to-barb adapters (Darwin Microfluidics) were incorporated to establish robust syringe connections. Chips were maintained at 37°C during preparation using a portable incubator and operated at room temperature (24°C) during MR acquisitions.

For cell suspension studies, cells were loaded into the experimental wells in 200 μL of culture medium. Hyperpolarised [1-^13^C]pyruvate was mixed with fresh medium immediately after dissolution (see section 4.7) and delivered through the microfluidic inlet to achieve a final pyruvate concentration of 3.2 mM and a final volume of 300 μL per well. Following substrate delivery, an air chase was applied to minimise dead volume within the microfluidic channels and promote mixing within the wells.

For scaffold studies, cell-seeded cryogels (see section 4.4) supported by nylon meshes were positioned within the experimental wells before chip assembly. Hyperpolarised substrate was delivered by injecting 360 μL of 40 mM hyperpolarised [1-^13^C]pyruvate mixed with 4.14 mL of culture medium while excess medium was removed from the outlet, resulting in perfusion of all scaffold-containing wells.

### 4.10. Longitudinal experimental set-up

A schematic representation of the longitudinal experimental platform is shown in Figure 5a,b. Longitudinal studies were performed using HepG2-seeded cryogel scaffolds loaded into the 8wmc, with one scaffold positioned in each experimental well.

Following each HP-MR acquisition, residual pyruvate was removed by flushing the chip with fresh culture medium. The inlet and outlet tubing were subsequently connected to a syringe-pump-based recirculation system (Darwin Microfluidics), enabling continuous perfusion of fresh medium through the chip at 0.1 mL min^-1^ for approximately 90 min. Recirculation was performed inside a portable incubator maintained at 37°C without CO_2_ supplementation.

Prior to each hyperpolarised acquisition, air bubbles were removed from the experimental wells to ensure homogeneous substrate delivery. Each experiment consisted of five HP-MR measurements separated by four recirculation periods. Cell metabolic activity following the complete protocol was assessed using the alamarBlue HS assay.

### 4.11. HP-MR data acquisition

All experiments were performed on a 3 T MRI scanner (BioSpec, 105 mm bore, Bruker) controlled using ParaVision 360 v3.4. Data from cell suspension and scaffold experiments were acquired using a dual-tuned ^1^H/^13^C transmit/receive volume and surface RF coil (Bruker, Germany), respectively. Prior to each acquisition, standard global and localised shimming procedures were performed over the experimental region.

Chip localisation was performed using T_2_-weighted TurboRARE images with echo time of 60-77 ms, repetition time of 1000-2000 ms, a 90-degree flip angle, 1-2 averages, an image pixel size of 192 x 192 for the 8wmc and 300 x 300 for the 4wmc, a field of view (FOV) of 42 mm x 42 mm for the 8wmc and 40 x 15 mm for the 4wmc, a slice thickness of 20 mm for the 8wmc and 15 mm for the 4wmc, with one imaging slice, 1-2 dummy scans, a Rare encoding order, bandwidth of 24691.4 Hz, a working frequency of 127.6 MHz and a working chemical shift of 4.7 ppm.

For single-time-point acquisitions, the hyperpolarised substrate was injected outside the scanner immediately before positioning the chip within the magnet bore. A single chemical shift imaging (CSI) dataset was acquired 35 s after dissolution using the CSI pulse sequence with a 40 mm x 15 mm FOV for the 4wmc and a 42-45 mm x 22-25 mm FOV for the 8wmc, an 8-10x16 voxel grid size, a 0.8-1.24 ms echo time, a 321.4-323.2 ms repetition time, a 22-degree flip angle, gaussian or block pulse shapes, a centric encoding order and compress sensing with 50% sampling and a 25% fully sampled center area.

Time-resolved experiments were performed using a sequential single-pulse slice-selective (SSPSS) acquisition scheme (Figure 4a). These experiments were performed using the 4wmc, with HepG2 cells loaded into three wells and cell-free medium loaded into the remaining well. Twenty time points were acquired per well following substrate injection with a 0.5 ms echo time, a 318.3 ms repetition time and 22-degree flip angle block pulses.

Longitudinal scaffold experiments were performed using the 8wmc and a dual-tuned ^1H/^13C surface coil. Two slice-selective regions encompassing four scaffolds each were acquired 35 s after substrate delivery using a 0.74 ms echo time, a 319.9 ms repetition time and 90-degree flip angle block pulses.

For all experiments using 13C spectroscopy, we used a 3236.3 Hz bandwidth, a 1.6 Hz/points spectral resolution, a 32.1 MHz working frequency and a 170 ppm working chemical shift.

### 4.12. CSI data processing

CSI datasets were reconstructed and analysed using custom MATLAB software (https://github.com/MIPMED-lab/CheShImP_DavidMIPMED) and custom Python scripts. Raw spectroscopic data were reconstructed in magnitude mode and apodised before analysis. Experimental wells were identified using the corresponding ^1^H-MRI images. Voxels containing noise only or substantial signal contamination from neighbouring regions were excluded from further analysis.

For each well, spectra from the selected voxels were summed prior to quantification. Metabolic activity was quantified as the lactate-to-pyruvate integral ratio (Lac/Pyr), and signal-to-noise ratio (SNR) was calculated from the summed spectra.

### 4.13. SSPSS data processing

Sequential acquisitions were processed using custom Python software. Raw free induction decays were apodised, Fourier transformed and phase corrected before spectral quantification. Acquisition metadata were used to automatically assign spectra to individual wells and acquisition times.

Lactate and pyruvate dynamics were analysed using a two-site exchange model described in the Supplementary Information to estimate apparent longitudinal relaxation times and exchange rate constants (k_PL_ and k_LP_). Peak integration ranges were manually defined. Longitudinal scaffold datasets were processed using the same workflow, although kinetic fitting was not performed because only a single post-injection time point was acquired.

### 4.14. Statistical analyses

Statistical analyses were performed using custom Python scripts. Pairwise comparisons were evaluated using two-tailed Student’s t-tests. Statistical significance was defined as *P* < 0.05.

### 4.15. Reporting summary

Further information on research design is available in the Nature Portfolio Reporting Summary linked to this article.

## 5. DATA AVAILABILITY

All raw, processed and analysed experimental data and engineering drawings associated with this study are publicly available through the CORA repository at doi.org/10.34810/data3242. These datasets include, but are not limited to, the source data underlying all figures and supplementary figures presented in this manuscript.

## 6. CODE AVAILABILITY

All custom code generated for this study is publicly available through the GitHub repository “BioWells HP-NMR” (https://github.com/MIPMED-lab/BioWells_HP-NMR). The source code developed for chemical shift imaging processing, visualisation and analysis is available through the GitHub repository “CheShImP” (https://github.com/MIPMED-lab/CheShImP_DavidMIPMED).

## 7. AUTHOR CONTRIBUTIONS

D.G.C. and I.M.R. designed experimentation. D.G.C., J.E. and M.A. developed all the spectroscopy, image and spectroscopic image protocols and acquisitions. L.M.F., A.H.G. and D.G.C. performed cell culturing and preparation for all the experiments. D.G.C., G.M and M.A. handled the dDNP machine for all experiments. G.M. build microfluidic chips, stop meshes and 3D printable designs. D.G.C. performed all data analysis and script development. J.E. and M.A. aided in the development and integrity of CheShImP. D.G.C., L.M.F., G.M. and I.M.R. wrote the manuscript. All the authors reviewed the manuscript. I.M.R. conceived and conceptualised the project, provided guidance and raised funding.

## 9. ACKNOWLEDGEMENTS

The authors would like to thank William Mander for his assistance and technical support with the HyperSense polariser; José Yeste and Maria Alejandra Ortega Machuca for discussions on microfludic chip and experimental design; Alejandro Portela for his input on the stop-mesh structures; Peter Sperling for his assistance with cell cultures; and the MicroFabSpace and Microscopy Characterisation Facility, Unit 7 of ICTS “NANBIOSIS” from CIBER-BBN at IBEC.

This work has received funding from: the European Union (GA-101195272, Q-AID) and (GA-101291716, CAMP), a European Union ERC Starting Grant (GA-101165045, LIFETIME); the Spanish grants with reference PID2023-151470OB-I00 funded by MICIU/AEI/ 10.13039/501100011033 and by “ERDF/EU” (METACHIP), RYC2020-029099-I funded by MCIN/AEI10.13039/501100011033 and by “ESF Investing in your future”, PLEC2022-009256 funded by MCIN/AEI/10.13039/501100011033 and by the European Union NextGenerationEU/PRTR (FLASHonCHIP), a grant within the framework Biotechnology Plan Applied to Health funded by MCIN/AEI10.13039/501100011033 and co-financed by the Spanish Ministry of Science and Innovation with funds from the European Union NextGenerationEU from the Recovery, Transformation and Resilience Plan (PRTR-C17.I1); from the Autonomous Community of Catalonia within the framework of the Biotechnology Plan Applied to Health; the BIST (Barcelona Institute of Science and Technology)-“la Caixa” Banking Foundation Chemical Biology programme. *Views and opinions expressed are however those of the author only and do not necessarily reflect those of the European Union or the European Research Council. Neither the European Union nor the granting authority can be held responsible for them*.

