## Supplementary Data for "A deployable microfluidic platform for parallel and longitudinal hyperpolarised magnetic resonance metabolic phenotyping"

### A Mathematical Model for Kinetic Rates

In this work, we use a common mathematical model used in the hyperpolarised NMR field [1–4]. The model describes the conversion from pyruvate (Pyr) to lactate (Lac) at constant rate $k_{PL}$, and its reversible reaction $k_{LP}$. Both hyperpolarised states are subject to longitudinal relaxation observability exponential decays described by the $T_1$s. In all cases, both pyruvate and lactate are labelled in the first carbon with a $^{13}C$ isotope.

The reaction:

$$\mathrm{Pyr} \underset{k_{LP}}{\overset{k_{PL}}{\rightleftharpoons}} \mathrm{Lac}$$
$$\downarrow T_1^P \qquad \downarrow T_1^L$$

The derived mathematical model following mass action kinetics:

$$\begin{aligned}\frac{d[Pyr]}{dt} &= k_{LP}[Lac] - (\frac{1}{T_1^P} + k_{PL})[Pyr] \\ \frac{d[Lac]}{dt} &= k_{PL}[Pyr] - (\frac{1}{T_1^L} + k_{LP})[Lac]\end{aligned} \tag{1}$$

#### B Supplementary Figures

This section includes all figures including relevant data for the manuscript that where not included in the main text.

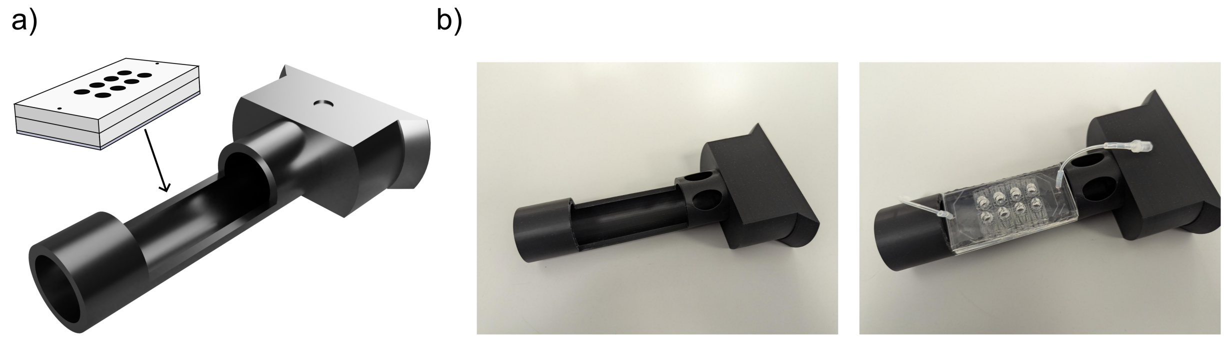


**Suppl. Fig. 1**: 3-D custom design microfluidic chip holder insert for a 3T Bruker Biospec MRI scanner with a dual-tuned $^{1}H$-$^{13}C$ volume coil (42 mm inner diameter). **a,** Schematic of the holder insert and the positioning of the microfluidic chips. **b,** Pictures of the custom-designed holder insert without (left) and with (right) a microfluidic chip.

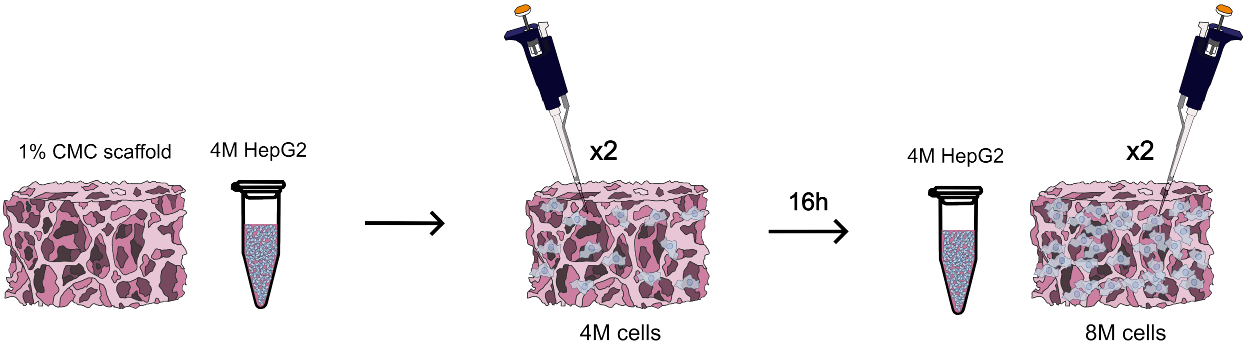


**Suppl. Fig. 2**: HepG2 cell seeding in the CMC scaffold. First, 4M cells were seeded in the CMC scaffold divided in 2 consequtive seedings of 2M cells separed by 20 minutes. The next day, we repeated the process to obtain a total of 8M cells per scaffold.

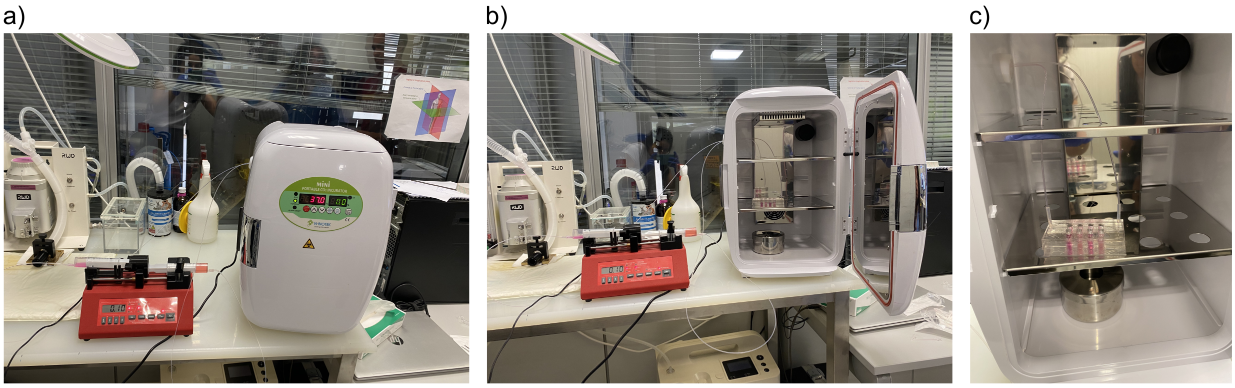

**Suppl. Fig. 3**: Experimental set up for longitudinal metabolic imaging experiments with cell seeded scaffolds. **a,** Picture of the portable incubator and syringe pump used for longitudinal experiments. **b,** Same as panel **b**, but with the incubator door opened to visualise the positioning and tubing connection of the microfluidic chip. **c,** Close up picture of the microfluidic chip containing four cell seeded scaffolds and recirculating media thanks to the microfluidics set up.

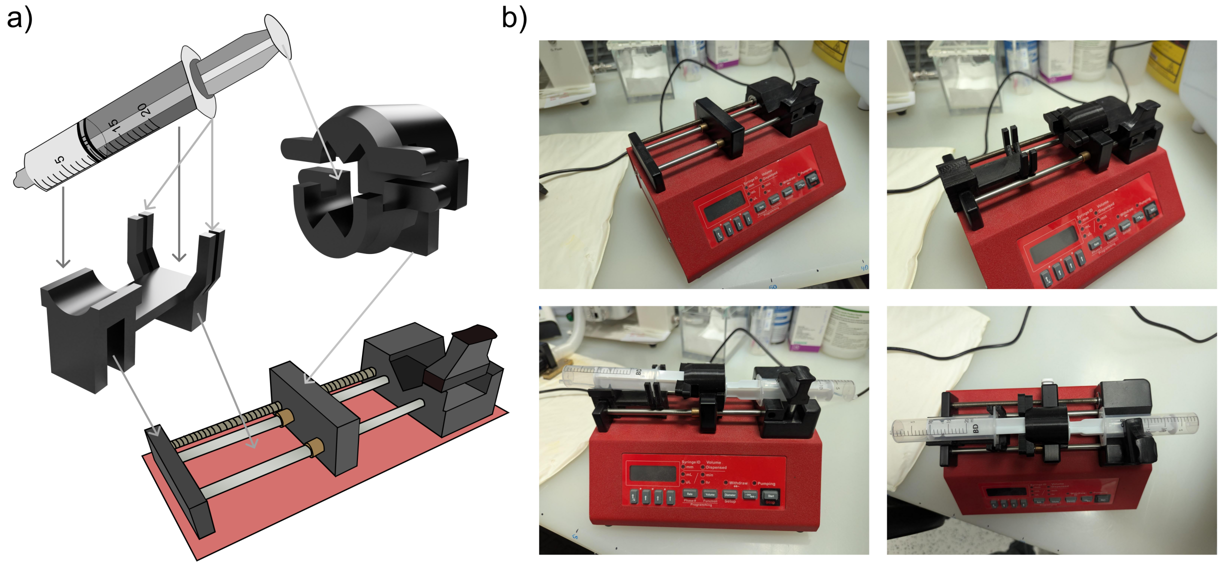

**Suppl. Fig. 4**: 3D custom-designed syringe support for simultaneous media suction and pumping out for microfluidic systems using a commercial pressure pump. **a,** Schematic for the syringe holder components indicating the placement of the syringe (20 mL) into this, and its placement into the syringe pressure pump. **b,** Pictures of the NE-300 InfusionONE syringe pump used in this work (top left), the same with the syringe holder components attached to it (top right) and the final set-up with the two syringes attached (bottom left and right).

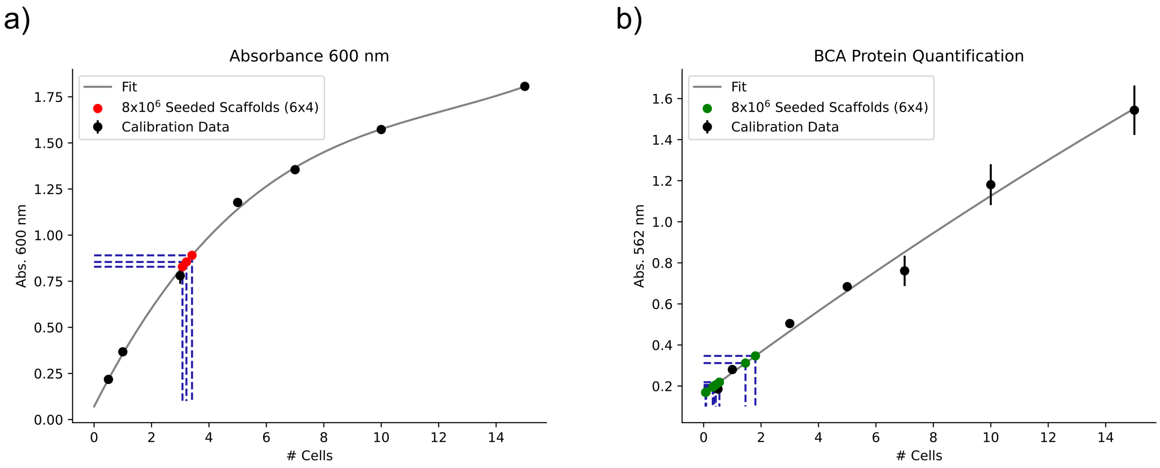


**Suppl. Fig. 5**: Cell number estimation within the cryogel scaffolds after seeding of eight million. **a,** Optical density analysis results to estimate cell number. **b,** BCA (Bicinchoninic Acid) protein quantification colorimetric assay test to estimate cell number. For all tests, we used media containing empty scaffolds as blanks.

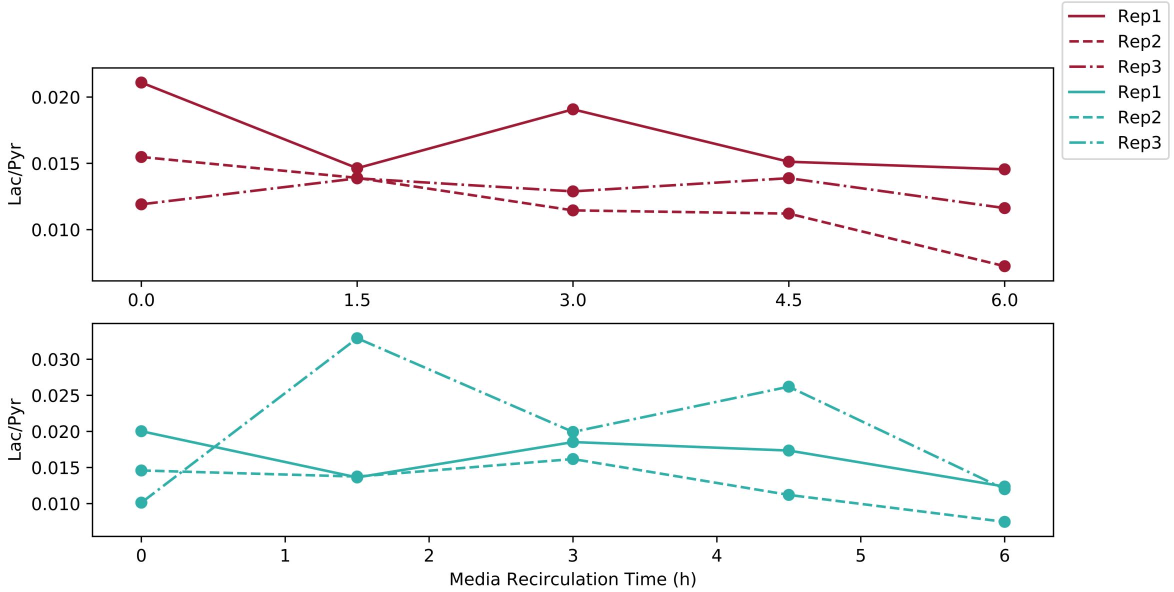


**Suppl. Fig. 6**: Lac/Pyr ratio evolution of each chip individually as a function of incubation time. Left region of every chip shown in red and right region shown in blue. Different line types to differenciate each chip.

| Figure | dDNP shots | Conditions per shot | HP-MR measurements | Conventional dDNP shots required |
|---|---|---|---|---|
| Fig. 2a | 5 | 8 | 40 | 40 |
| Fig. 2b | 5 | 8 | 40 | 40 |
| Fig. 3a | 5 | 4 | 20 | 20 |
| Fig. 3b | 3 | 4 | 12 | 12 |
| Fig. 3c | 3 | 4 | 12 | 12 |
| Fig. 4 | 8 | 4 | 32 | 32 |
| Fig.5 (longitudinal, 3 replicates) | 15 | 2 | 30 | 30 |
| **Total** | 44 | - | 186 | 186 |

| Metric | Value |
|---|---|
| Total dDNP events used | **44** |
| Equivalent conventional dDNP events | **186** |
| Increase in HP-MR throughput | **4.2x** |
| Increase in longitudinal experiments throughput | **5x** |
| Experimental cost saved (€200/event) | **€28,400** |

**Suppl. Table 1**: Quantification of throughput gains achieved by the parallel HP-MR platform

.
